# Gelsolin protects mitochondria and regulates inflammation during *Legionella pneumophila* infection

**DOI:** 10.64898/2026.08.17.745205

**Authors:** Owen D. Whitham, Mostafa M. Eltobgy, Mohamed M. Shamseldin, Asmaa Badr, Richard A. Perez, Yara Y. Hassan, Heba M. Amer, Gauruv Gupta, Sarah E. Faber, Frank H. Robledo-Avila, Xiaoli Zhang, Maya Mikami, Santiago Partida-Sanchez, Stephanie Seveau, Amal O. Amer

## Abstract

*Legionella pneumophila* (*L. pneumophila*) is the causative agent of Legionnaires’ disease, a severe bacterial pneumonia. Difficulty in diagnosing Legionnaires’ disease leads to an underreporting of cases and delayed treatment. Rapid-acting, broad-spectrum therapies are needed to treat pathology while avoiding antibiotic resistance. We showed that gelsolin knockout (*gsn*^-/-^) mice succumb more quickly to severe *L. pneumophila* infection despite no difference in bacterial loads in the lung compared to wild type mice. There is an increase in CXCL1/KC production from macrophages from *gsn*^-/-^ mice, which is accompanied by increased neutrophils and apoptosis in their lungs. Neutrophils lacking gelsolin produce fewer neutrophil extracellular traps, and their mitochondrial capacity is diminished in response to *L. pneumophila*. Gelsolin is required for maintaining mitochondrial network morphology and respiration in *L. pneumophila* infected macrophages. When given recombinant gelsolin protein, *gsn*^-/-^ mice survive significantly longer during severe *L. pneumophila* infection, with reduced lung pathology, and the inflammatory signature of their macrophages was reduced *in vitro*. Together, gelsolin protects mice during severe *L. pneumophila* infection, dampens inflammation, promotes mitochondrial health, and maintains neutrophil function.

## Introduction

*Legionella pneumophila* (*L. pneumophila*) is a gram-negative respiratory intracellular pathogen that is the causative agent of the severe atypical pneumonia Legionnaires’ disease. Contamination of domestic or industrial water systems, such as air conditioning systems or water-cooling towers, and formation of biofilms can lead to aerosolization and subsequent inhalation of *L. pneumophila* (De Filippis et al., 2018; Fang et al., 2023; Van Heijnsbergen et al., 2015). *L. pneumophila* is phagocytosed by alveolar macrophages and secretes effector proteins to halt phagosome-lysosome fusion, creating the *L. pneumophila* containing vacuole (LCV) (Horwitz, 1983). During a successful infection cycle, *L. pneumophila* replicates and lyses the LCV, allowing the infectious progeny to infect a new cell. However, *L. pneumophila*-derived pathogen-associated molecular patterns, such as flagellin or lipopolysaccharide, escape from the LCV and are detected by the host’s pattern recognition receptors, specifically Naip5/NLRC4 or caspase-4/11, which promote the activation of the canonical and non-canonical inflammasomes respectively (Amer et al., 2006b; Caution et al., 2015b; Cerqueira et al., 2015; Luo, 2012; Pereira, Marques, et al., 2011). In both cases, inflammasome activation leads to pyroptotic cell death and inflammatory burst that limits the spread of the pathogen (Cerqueira et al., 2015; Pereira, Marques, et al., 2011; Pereira, Morgantetti, et al., 2011; Zhao et al., 2011). Pathogen clearance and regulation of the inflammatory response are essential to the severity of Legionnaires disease. Individuals who are elderly, immunodeficient, or have a history of smoking are at an increased risk of experiencing severe disease (Dartevel et al., 2025; Lupia et al., 2023). Early diagnosis and rapid treatment are critical to limit disease progression and prevent poor outcomes for these individuals.

Gelsolin (GSN) is a calcium-dependent actin severing and capping protein with two isoforms, plasma (pGSN) and cellular gelsolin (cGSN), that arise through alternative splicing of a single *gsn* gene (Bryan, 1988; Stella et al., 1994). cGSN regulates cellular processes including actin dynamics, apoptosis, and mitochondrial homeostasis (García-Bartolomé et al., 2017; Geng et al., 1998; Janmey et al., 1985; Kothakota et al., 1997a; Koya et al., 2000; Kusano et al., 2000). pGSN contains a 25 amino acid N-terminal secretion signal that facilitates its secretion, primarily from muscle cells, into the blood, where it contributes to the clearance of circulating actin and inactivation of inflammatory molecules (Bucki et al., 2005, 2010, 2014; Kwiatkowski et al., 1988; Lind et al., 1986). In humans, decreased pGSN (and increased F-actin) concentrations in the blood correlate with worsened disease severity and outcome in patients with a broad range of infections, injuries, and diseases (DiNubile et al., 2002; Esawy et al., 2020; P. S. Lee et al., 2006, 2008; Self et al., 2019; Shi et al., 2014). Gelsolin is integral to the immune function of neutrophils and macrophages, and recombinant human gelsolin (rGSN) has been shown to reduce neutrophil accumulation in the lungs in a mouse model of acute hyperoxia (Christofidou-Solomidou et al., 2002; Fettucciari et al., 2015; J. Lee et al., 2024; Mannherz et al., 2024a; Serrander et al., 2000). Several studies have demonstrated the protective effects of rGSN administration in animal models of infection, but none have done so in *L. pneumophila* infected mice (Cheng et al., 2017; Cui et al., 2022; DiNubile et al., 2020; Kobzik et al., 2020; Piktel et al., 2020; Yang et al., 2015). While it is accepted that gelsolin deficiency aggravates pathology in a variety of infectious diseases, few of these studies have investigated this directly using a global gelsolin knockout mouse model, and none have done so in a mouse model of *L. pneumophila* infection.

Human caspase-4 and its murine ortholog caspase-11 mediate non-canonical inflammasome activity, a central innate immune pathway in the response to many intracellular pathogens including *L. pneumophila* (Akhter et al., 2012a; Kayagaki et al., 2011). The activation of caspase-4/11 or caspase-1, through non-canonical and canonical inflammasome activation respectively, coordinates a highly inflammatory cell death termed pyroptosis that is vital for the control of such pathogens (Cerqueira et al., 2015; Kayagaki et al., 2015). While the role of caspase-1 has been studied in detail in many diseases, novel functions and substrates of caspase-4/11 continue to be discovered and remain incompletely understood. We have previously shown that caspase-4/11 mediates several cellular processes including actin organization and mitochondrial function during infection (Akhter et al., 2012a; Caution et al., 2015a; Krause et al., 2019). Here we establish gelsolin as a possible substrate candidate of noncanonical inflammasome activation, given its susceptibility to cleavage by caspase-4, and demonstrate its role in regulating Legionnaires disease pathology and severity.

## Results

### Gelsolin deficient mice rapidly succumb to severe *L. pneumophila* infection

Several studies have shown that plasma gelsolin is protective in several animal models of infection and inflammation, and pGSN concentrations negatively correlate with disease severity in critically ill patients (Cheng et al., 2017; Cui et al., 2022; DiNubile et al., 2002, 2020; Esawy et al., 2020; Kobzik et al., 2020; P. S. Lee et al., 2006, 2007, 2008; Piktel et al., 2020; Self et al., 2019; Shi et al., 2014; Yang et al., 2015). To determine if gelsolin is protective during severe *L. pneumophila* infection, we infected wild type (WT) and gelsolin knockout (*gsn*^-/-^) mice with a severe dose (5x10^6^ CFU) of *L. pneumophila*. *Gsn*^-/-^ mice succumbed to severe infection more rapidly compared to WT mice (Fig 1A). Neither WT nor *gsn*^-/-^ mice infected with a physiological dose of *L. pneumophila* succumbed to infection (1x10^6^ CFU) (Data not shown). This indicates that gelsolin is vital for survival of the host during severe *L. pneumophila* infection, but not during mild infection. The lung function of mice infected with the severe dose of *L. pneumophila* was assessed by whole body plethysmography (WBP), a non-invasive method for quantifying lung function and respiratory distress. WBP analysis showed that WT mice had a significantly higher airway resistance as measured by enhanced pause (Penh) at early stages of the infection (Fig 1B). However, the airway constriction in WT mice respiration steadily recovered over time, while *gsn*^-/-^ mice failed to recover (Fig 1C). This indicates that gelsolin affects lung functions in response to *L. pneumophila* infection and is also important in restoring the lung to its healthy state.

**Figure 1.**
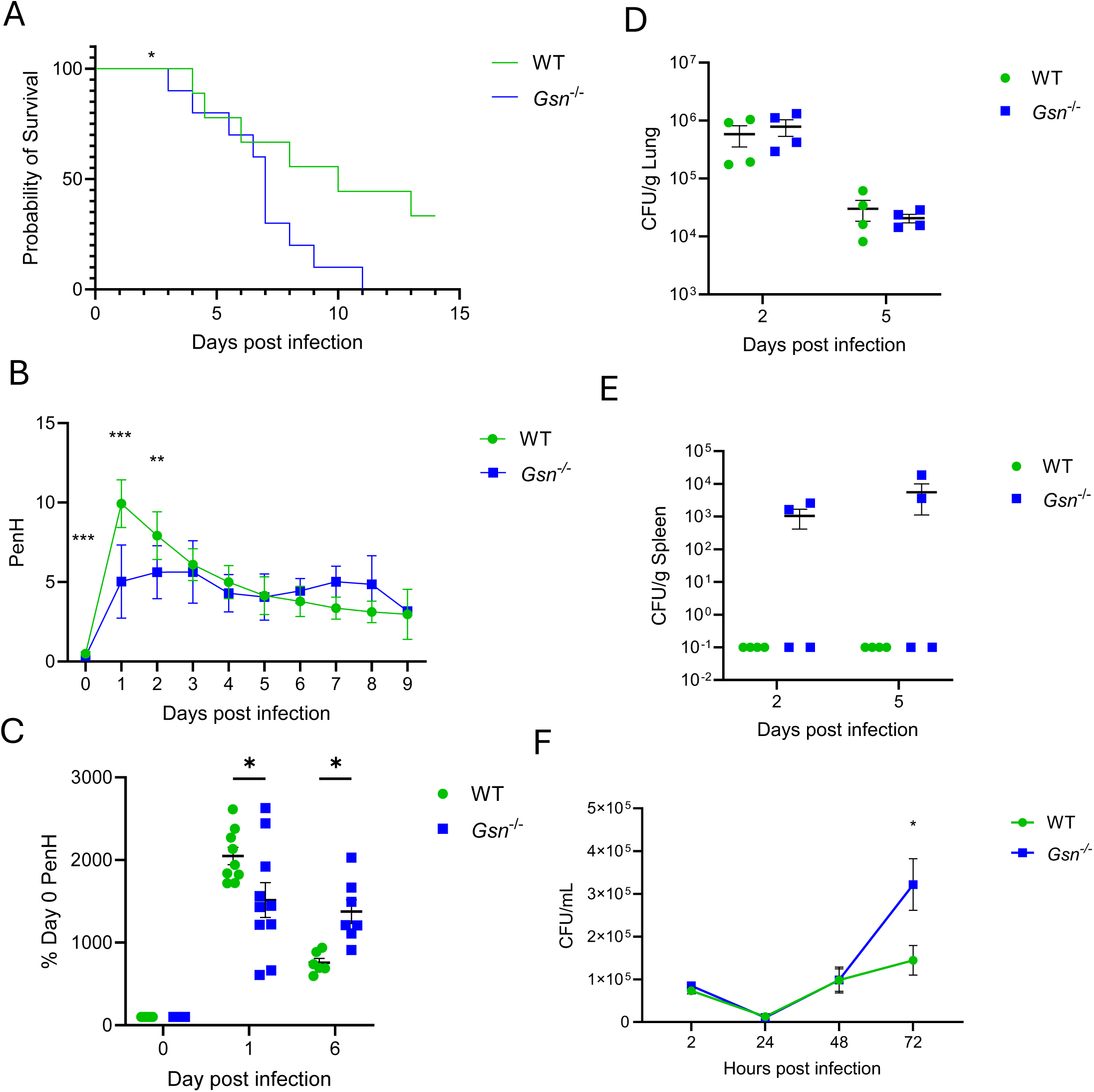
Gelsolin protects against lethal *L. pneumophila* infection *in vivo*. (A) Kaplan-Meier survival curve and (B) lung function (unpaired t-test) measured by whole body plethysmography of WT (n=9) and *gsn*^-/-^ (n-10) mice infected with a lethal dose (5x10^6^ CFU) of LP02. Two-sample t-test with multiple comparison adjustment. (C) Penh from *B* as a percent increase from baseline D0. Two-sample t-test with multiple comparison adjustment. (D,E) CFUs recovered from (D) lung and (E) spleen of WT (n=4) and *gsn^-^*^/-^ (n=4) mice infected with lethal dose of LP02 at day 2 or day 5 post infection. Two-sample t-test with multiple comparison adjustment. (F) Enumeration of *Legionella* colony forming units from WT and *gsn*^-/-^ macrophages (n=5) infected with an MOI of 0.025 LP02W at 2, 24, 48, and 72h post infection. Two-sample t-test with multiple comparison adjustment. Error bars represent SEM. *p<0.05, **p<0.01, ***p<0.001.

To assess if the reduced survival of the *gsn*^-/-^ mice was due to increased bacterial growth, bacterial burden in the lung and spleen of WT and *gsn*^-/-^ mice were quantified at 2- and 5-days post infection. We found no difference in bacterial burden in the lung between WT and *gsn*^-/-^ mice (Fig 1D). When we quantified bacterial burden in the spleens of these mice, we found more *gsn*^-/-^ mice with *L. pneumophila* in the spleen compared to WT, indicating an increased susceptibility to bacterial dissemination (Fig 1E). Interestingly, *in vitro*, the *L. pneumophila* burden in *gsn*^-/-^ macrophages after 72h infection was significantly higher than that of the WT (Fig 1F). This indicates that while gelsolin is required for *L. pneumophila* clearance in the macrophage, the complexity of the lung environment and the contributions of other cells introduce factors that might mask this deficit.

### Gelsolin deficiency promotes neutrophil accumulation in the lungs

Gelsolin is a potent actin regulating protein and has been shown to mediate cellular motility and migration of numerous cell types, including neutrophils (Christofidou-Solomidou et al., 2002). Like other pneumonias, Legionnaires’ disease pathology is partially mediated by an inflammatory response and influx of immune cells, resulting in lung injury. However, an incompetent immune cell response in the lung can lead to prolonged infection. Due to the higher degree of lung distress but higher survival in WT mice compared to *gsn*^-/-^, we assessed the inflammation in the lungs of these mice. First, we measured the inflammatory cytokines released by *L. pneumophila* infected macrophages. While infected *gsn*^-/-^macrophages had more CXCL1/KC (KC) (Fig 2A), a potent neutrophil-recruiting chemokine, and IL-6 (Fig 2B) than WT mice, IL-1β (Fig 2C) and TNFα (Fig 2D) were not elevated, indicating specificity in the augmentation of inflammatory cytokines by endogenous gelsolin. We infected mice with severe dose of *L. pneumophila* and quantified cytokines in the lungs at 2- and 5-days post infection. We performed H&E staining on the lungs of the same mice and there was no difference in gross lung pathology or cellularity (Fig 2E). However, staining for fragmented DNA by Terminal deoxynucleotidyl transferase dUTP nick end labeling (TUNEL) showed a significantly higher signature of apoptosis in almost all cells in the *gsn*^-/-^ lung compared to WT at day 2 (Fig 2F). Immunohistochemistry staining for CD45+ immune cells shows a unique accumulation of infiltrating immune cells in *gsn*^-/-^ lungs that seemingly fail to penetrate the alveolar space (Fig 2G). When measuring inflammatory cells in the lungs by flow cytometry, we found that lung infiltrating neutrophils (CD45^-^ CD11b^+^ Ly6G^+^) were increased in the lungs of *gsn*^-/-^ mice, correlating with the increase in macrophage-derived KC (Fig 2H). This indicates that gelsolin is required for limiting KC production by macrophages and reducing infiltration of neutrophils into the lung during severe *L. pneumophila* infection.

**Figure 2.**
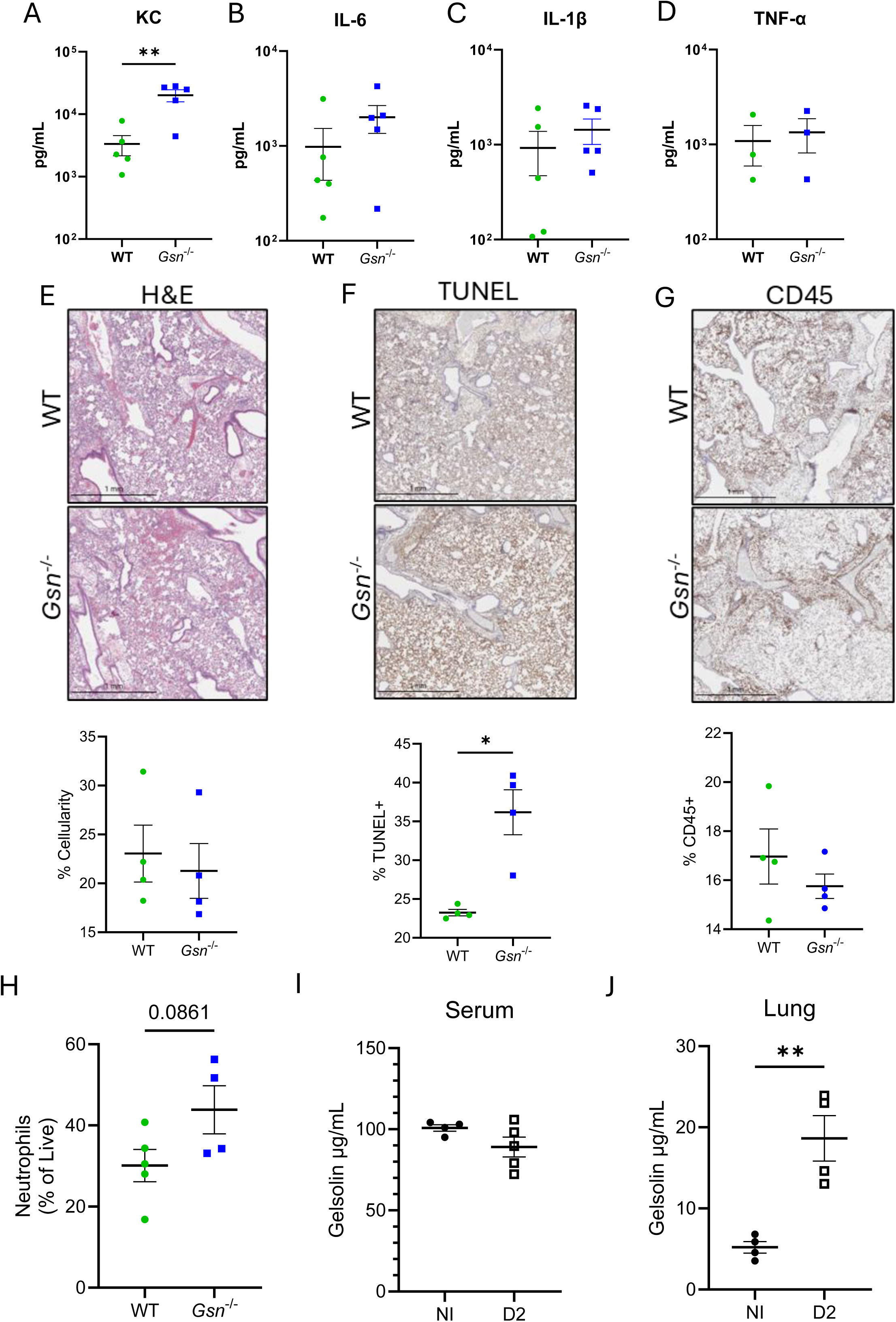
Gelsolin regulates inflammation in the lung during *L. pneumophila* infection. (A-D) CXCL1/KC (A), IL-6 (B), IL-1β (C), and TNF-α (D) measured by ELISA in the supernatant of WT (n=5) and *gsn^-^*^/-^ (n=5) macrophages infected with LP02 MOI 0.5 for 24 hours; unpaired t-test. (E-G) Representative histology and immunohistochemistry images of lung sections of WT (n=4) and *gsn*^-/-^ (n=4) control mice or mice infected with a lethal dose of LP02 for two days. *S*taining was performed with (E) hematoxylin & eosin (E), TUNEL (F), or with antibodies against CD45 (G) with quantification represented below; unpaired t-test. (H) Flow cytometry quantification of lung infiltrating neutrophils (CD45^-^ CD11b^+^ Ly6G^+^) in the lung of WT (n=5) and *gsn*^-/-^ (n=4) mice infected with lethal dose of LP02 at day 2 post infection; unpaired t-test. (I,J) Mouse gelsolin concentration, measured by ELISA in the (I) serum (I) and lung homogenate (F) of WT mice infected with LP02 (n=5) for two days or non-infected controls (n=4); unpaired t-test. Error bars represent SEM. *p<0.05, **p<0.01.

### Recombinant human gelsolin administration reduces *L. pneumophila* induced lung pathology and lethality

In humans, it is thought that pGSN is consumed at the site of injury, and as a result its systemic actin-clearing and anti-inflammatory functions are diminished. Similarly, we observed that GSN decreased in the serum of *L. pneumophila* infected mice, while it accumulated in their lungs (Fig 2I,J). Several studies have shown that recombinant human gelsolin (rGSN) improves the outcome of mice in models of sepsis, infection, inflammation, and lung injury (Cheng et al., 2017; DiNubile et al., 2020; Kobzik et al., 2020; Koya et al., 2000; Piktel et al., 2020). Administration of rGSN has been shown to reduce pathogenic neutrophil influx into the lung (Christofidou-Solomidou et al., 2002). To determine if the protective effects of gelsolin are mediated by pGSN or cGSN, WT and *gsn*^-/-^ mice were infected with a high dose of *L. pneumophila*. Immediately after infection, mice were injected intraperitoneally with either PBS (WT and *gsn*^-/-^ mice) or 350µg rGSN (*gsn*^-/-^ mice). These mice were treated with PBS or rGSN daily by intraperitoneal injection (IP), monitored for weight and survival, and lung function was assessed daily by WBP. We found that *gsn*^-/-^mice receiving rGSN survived significantly longer than those receiving PBS (Fig 3A). Interestingly, airway constriction of *gsn*^-/-^ mice receiving rGSN was significantly lower than that of the WT and *gsn*^-/-^ mice receiving PBS (Fig 3B). rGSN had no profound effect in inflammatory cytokines in the serum or lungs. Lung pathology at two days post infection in mice receiving rGSN was significantly reduced compared to mice receiving PBS (Fig 3C,D). However, there was no decrease in neutrophil accumulation in the lungs of mice receiving rGSN (Fig 3E). rGSN was detectable in the serum of naïve mice 24 hours after one injection (Fig S1A,B), and at an even higher concentration in the lungs and serum of *L. pneumophila* infected mice that received daily injections (Fig 3F,G). *In vitro*, addition of rGSN to *gsn*^-/-^ macrophages infected with *L. pneumophila* significantly decreased concentrations of IL-1β, KC, IL-6, and TNFα in the supernatant after 24 hours of infection (Fig 3H-K), highlighting the ability for exogenous gelsolin to dampen inflammation in a non-specific manner. Importantly, this reduction of cytokines was observed when rGSN was added after infection, indicating that gelsolin can suppress macrophage-derived cytokines independent of direct interactions with the bacteria.

**Figure 3.**
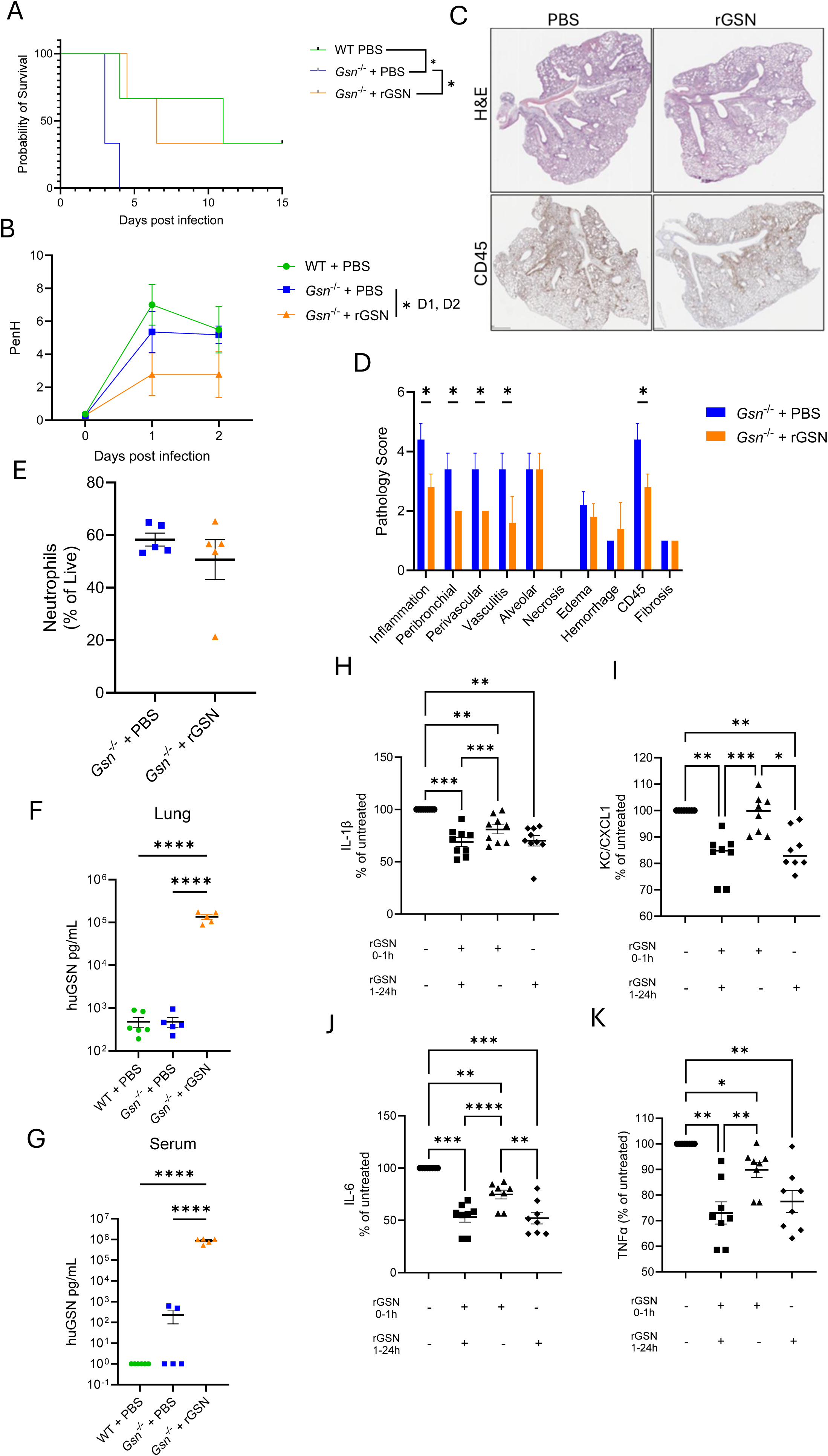
Recombinant human gelsolin reduces lethality and lung pathology during severe *L. pneumophila* infection. (A) Kaplan-Meier survival curve of WT (n=3) mice treated with PBS and *gsn*^-/-^mice treated with PBS (n=3) or rGSN (n=3) IP and infected with 5x10^6 LP02. (B) Airway resistance (PenH) as measured by whole body plethysmography in WT (n=6) mice treated with PBS and *gsn^-/-^*mice treated with PBS (n=5) or rGSN (n=5) infected with 5x10^6 colony forming units of LP02 and treated with PBS or rGSN for two days; unpaired t-test. (C) Representative images of lung H&E and CD45 staining from the same mice as in *B*. (D) Quantitative pathology scoring of images from C; Two-sample t-test with multiple comparison adjustment. (E) Flow cytometric quantification of lung infiltrating neutrophils (CD45^-^ CD11b^+^ Ly6G^+^) in the lungs of the same mice; unpaired t-test. (F,G) Human gelsolin measured by ELISA from the lung homogenate (F) and serum (G) from mice from B; unpaired t-test. (H-K) *Gsn*^-/-^macrophages were infected with LP02 MOI 0.5 for 24h and treated with PBS (-/-) or rGSN (100µg/mL) before (+/-), after (-/+), or before and after initial bacterial exposure (+/+) and IL-1β (H), CXCL1/KC (I), IL-6 (J), and TNFα (K) were measured in the supernatants by ELISA (n=8-9); one-way Anova with multiple comparisons. Error bars represent SEM. *p<0.05, **p<0.01, ***p<0.001, ****p<0.0001.

### Gelsolin is required for neutrophil NET formation and mitochondrial function in response to *L. pneumophila*

Given the increased expression of the neutrophil-attracting chemokine in the macrophages of *gsn*^-/-^ mice, as well as the increased neutrophils in the lung, we next investigated the role of gelsolin regarding intrinsic neutrophil function in response to *L. pneumophila* infection. It was previously reported that gelsolin contributes to NET formation by neutrophils stimulated with ionomycin (Mannherz et al., 2024). We isolated neutrophils from WT and *gsn*^-/-^ mice and treated them with *L. pneumophila,* PMA, or media, and measured NET formation by confocal microscopy. We found that *gsn*^-/-^ neutrophils produced significantly fewer NETs in all conditions (Fig 4A,B). Mitochondrial integrity and function are required for producing reactive oxygen species (ROS) and ROS-dependent NETs in response to pathogens (Douda et al., 2015). To determine if gelsolin regulates neutrophil respiration during *L. pneumophila* infection, we measured the mitochondrial respiration of WT and *gsn*^-/-^ neutrophils infected with *L. pneumophila* by seahorse assay using the mitochondrial stress test. We found that *gsn*^-/-^ neutrophils had a significantly lower spare respiratory capacity at rest and with *L. pneumophila* infection compared to WT neutrophils (Fig 4C). Therefore, gelsolin is required for NET formation and maintaining mitochondrial respiration in neutrophils during *L. pneumophila* infection.

**Figure 4.**
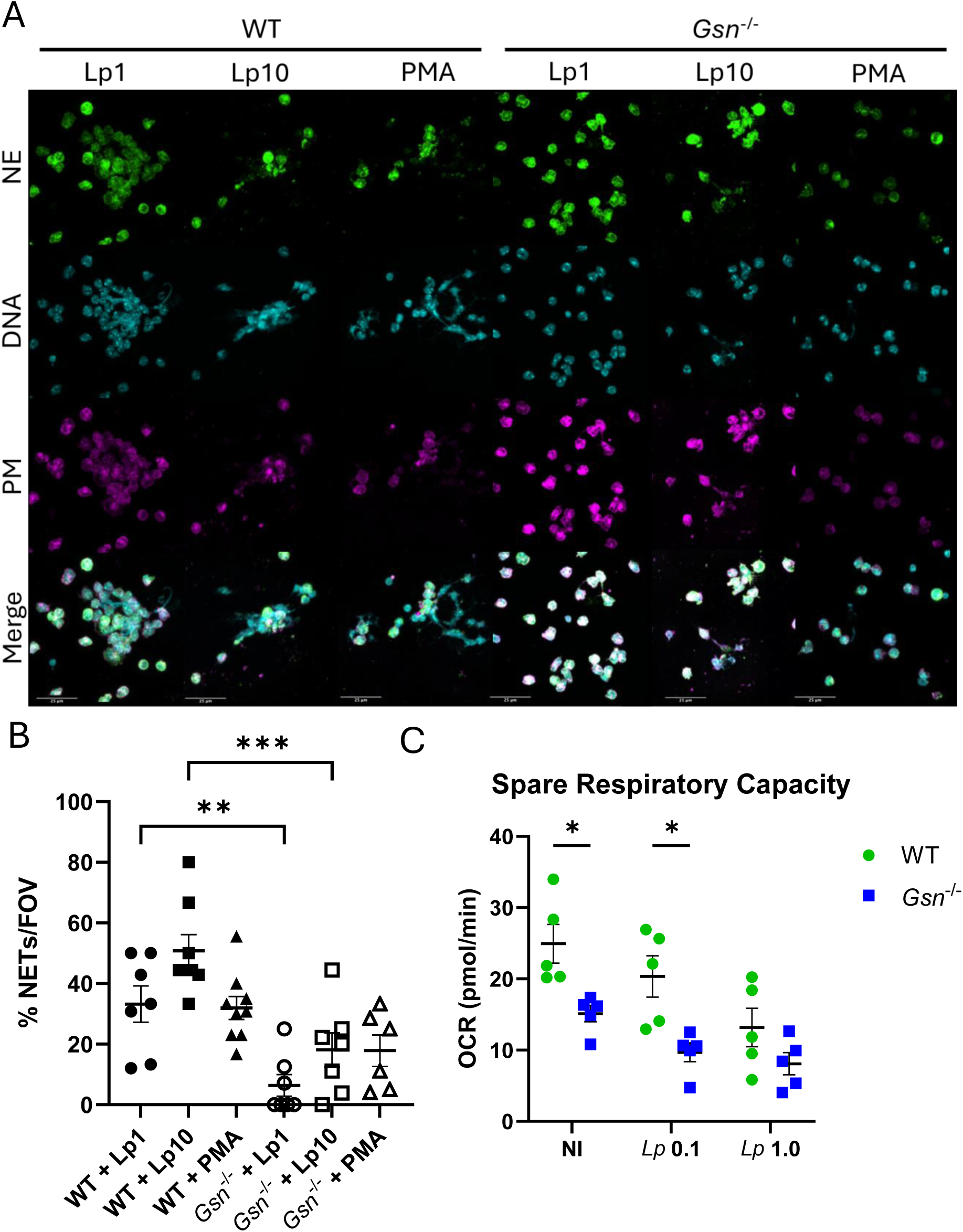
Gelsolin is required for NETosis and mitochondrial function in neutrophils in response to *L. pneumophila*. (A) WT (n=3) and *gsn*^-/-^ (n=3) neutrophils were treated with PMA (200nM) or LP02 (MOI 1 or 10) for 4 hours. Samples were stained for DNA, nuclear envelope, and plasma membrane and the NETs were imaged by confocal microscopy. Representative images from three independent experiments. (B) Quantification of the percentage of cells exuding NETs from A; one-way Anova with multiple comparisons. (C) Spare respiratory capacity of WT (n=5) and *gsn*^-/-^ (n=5) neutrophils measured by Seahorse mitochondrial stress test kit after infection with LP02 (MOIs 0.1 or 1) for 2 hours; Two-sample t-test with multiple comparison adjustment. Error bars represent SEM. *p<0.05, **p<0.01, ***p<0.001.

### Gelsolin maintains mitochondrial network and function during *L. pneumophila* infection

Gelsolin interacts directly with mitochondria and mediates mitochondrial functions including gating voltage dependent anion channel (VDAC) on the mitochondrial membrane (Kusano et al., 2000). During *L. pneumophila* infection, maintenance and augmentation of mitochondria is integral for pathogen defense and cellular function (Kajiwara et al., 2018). *L. pneumophila* can target the host mitochondria and shift metabolism to favor pathogen growth (Escoll et al., 2017; Fu et al., 2022). We investigated the role of gelsolin in maintaining mitochondrial network connectivity. Macrophages lacking gelsolin have a significantly higher frequency of mitochondrial network fracturing and rounding compared to WT macrophages (Fig 5A-C, supplemental movie S1 & S2), indicating a higher degree of stress and dysfunction. TEM imaging of these cells revealed fewer mitochondria near the bacterium in *gsn*^-/-^ macrophages compared to WT macrophages (5D). We analyzed the mitochondrial function of these macrophages by seahorse assay and observed a significantly decreased spare respiratory capacity in *gsn*^-/-^ macrophages compared to WT (5E). Therefore, gelsolin maintains mitochondrial network shape and overall mitochondrial function during *L. pneumophila* infection in macrophages.

**Figure 5.**
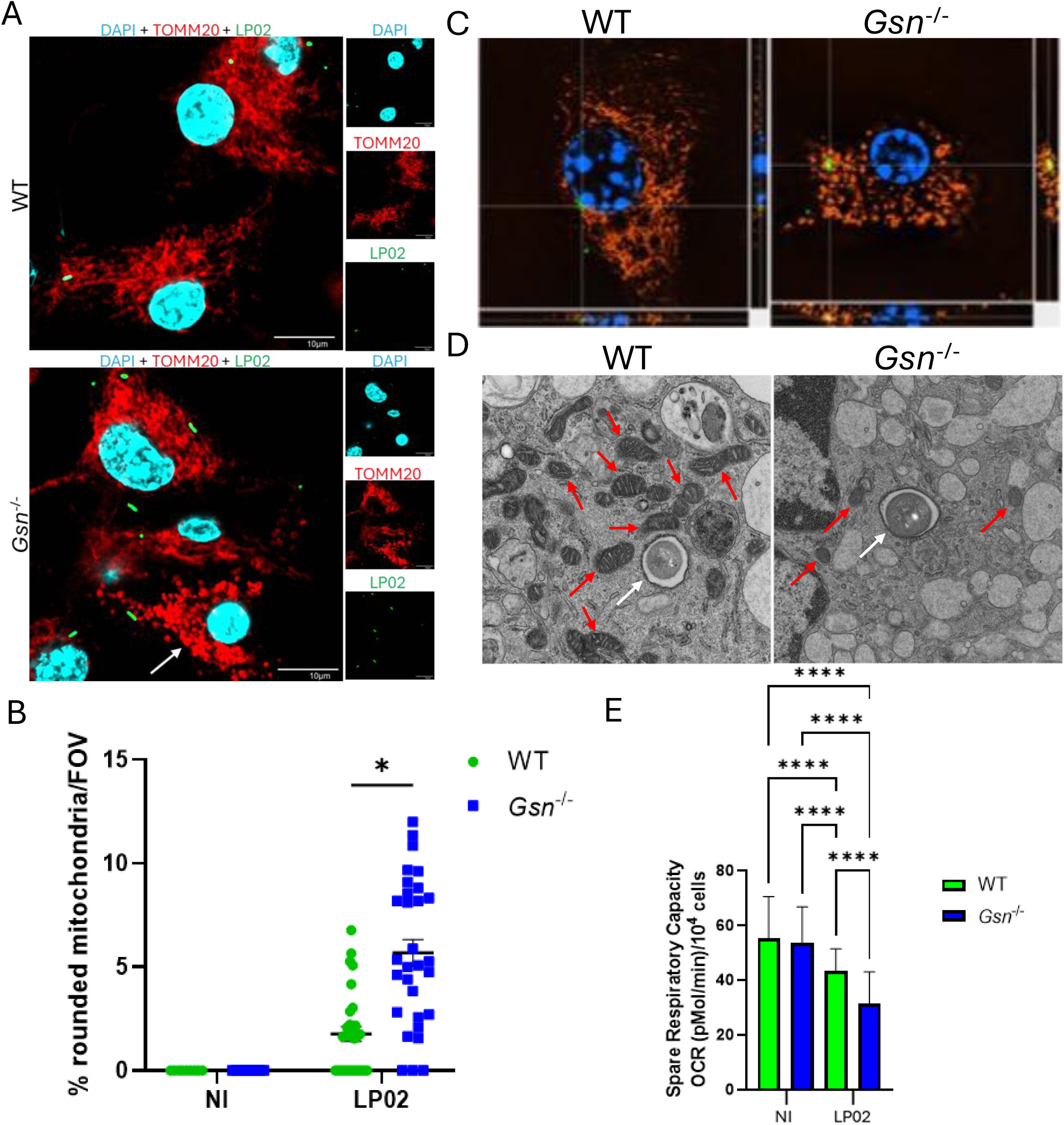
Gelsolin maintains mitochondrial shape and respiration during *L. pneumophila* infection. (A) Representative confocal microscopy images of WT and *gsn*^-/-^ macrophages (n=3) infected with LP02 MOI 0.5 for 1h and stained for TOMM20 (red) DAPI (cyan), and *L. pneumophila* (green). White arrow denotes cell with disrupted, rounded mitochondrial network. (B) Quantification of cells with disrupted, rounded mitochondrial networks from *A.* A total of over 1500 cells were counted for each condition over three independent replicates and 10 FOV per replicate; unpaired t-test. (C) SIM images of WT and *gsn*^-/-^macrophages infected with LP02 MOI 0.5 for 1h with the same staining from *A*, highlighting the disrupted mitochondrial morphology in *gsn*^-/-^ macrophages. (D) Representative TEM images of WT and *gsn*^-/-^ macrophages infected with LP02 MOI 0.5 for 1h. White arrows designate *Legionella* containing vacuole. Red arrows designate mitochondria in proximity to the bacteria. (E) Spare respiratory capacity normalized to cell count in WT or *gsn*^-/-^ macrophages infected with *L. pneumophila* for 1h followed by 1h 0% CO_2_ incubation; one-way Anova with multiple comparisons. Error bars represent SEM. *p<0.05, ****p<0.0001.

### Recombinant caspase-4 cleaves gelsolin into fragments distinct from caspase-3 but similar to caspase-1 generated products

Gelsolin is a canonical substrate of caspase-3, which once activated cleaves gelsolin at the S3-S4 linker to release the N-terminal half from the C-terminal (Kothakota et al., 1997a). The free N-terminal can sever actin in a calcium-independent manner and help execute the apoptotic functions of caspase-3 (Geng et al., 1998; Kothakota et al., 1997b). We have previously shown that caspase-11 activation alters actin dynamics in macrophages during *L. pneumophila* infection (Akhter et al., 2012a; Caution et al., 2015a). Due to the proteolytic nature of caspase-4/11 and gelsolin’s susceptibility to cleavage by other caspases, we sought to determine if human caspase-4 could cleave human recombinant gelsolin protein. We incubated recombinant human gelsolin with or without either recombinant human caspase-4, caspase-1, or caspase-3 protein at 37°C for 2 hours and reaction was stopped by the addition of protease inhibitor and incubation on ice. We analyzed the cleavage products by SDS-PAGE and western blot and found that caspase-4 produces a distinct cleavage signature than that of caspase-3, resulting in a higher number of smaller products (Fig 6). Due to the overlap of substrates and cleavage motifs between caspase-1 and caspase-4/11, we sought to determine if caspase-1 would generate gelsolin cleavage products like those generated by caspase-4. Indeed, caspase-1 produced a cleavage pattern with similar sized products to caspase-4 (Fig 6). We also incubated gelsolin with caspase-3 and caspase-4 simultaneously, or caspase-3 followed by caspase-4, and found that the GSN p39/41 caspase-3 products were completely gone, indicating that caspase-4 can further cleave these GSN fragments (Fig 6).

**Figure 6.**
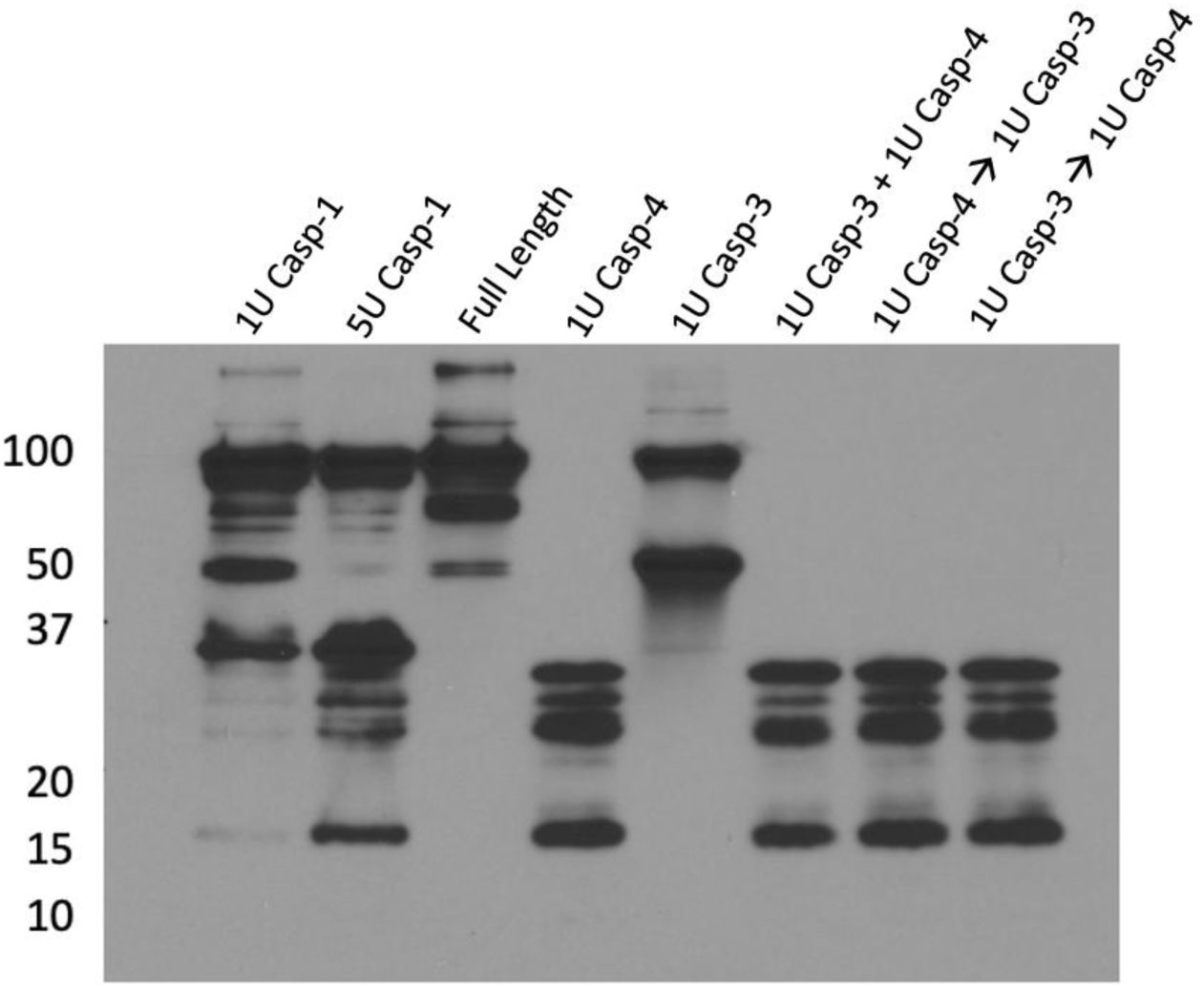
Gelsolin is cleaved by recombinant caspase-4, resulting in different fragments than produced by caspase-3 cleavage. Western blot of 1µg recombinant human gelsolin cleavage by 1 or 5 units of recombinant caspase-1, caspase-3, caspase-4, and combination of recombinant caspase-3 and caspase-4 simultaneously (+) or in succession (→) using anti-gelsolin antibody.

## Discussion

*L. pneumophila* causes the atypical community acquired pneumonia known as Legionnaires’ disease, a severe pneumonia that can be fatal in vulnerable populations (Dartevel et al., 2025; Lupia et al., 2023). Of particular concern is that *L. pneumophila* outbreaks are most prevalent in artificial water systems and buildings with complex water systems, including hospitals, retirement homes, and residential buildings, where vulnerable populations are exposed (De Filippis et al., 2018; Fang et al., 2023; Van Heijnsbergen et al., 2015). This necessitates the development of effective host-targeting treatments that are rapid-acting, broad spectrum, and non-immunogenic without contributing to antibiotic resistance. Additionally, *L. pneumophila* is a professional intracellular pathogen that alters cell biological pathways including actin machinery, metabolism, and inflammatory signaling to establish infection. To this effect, we show that rGSN administration improves the lung pathology of *L. pneumophila* infected mice at early timepoints, highlighting the potential of rGSN as a rapid-acting treatment for acutely infected patients. rGSN administration also improved the survival and lung function of infected mice, suggesting its potential to improve overall patient outcomes during severe Legionnaires’ disease.

Although rGSN administration did not contribute to bacterial clearance in our study, intravenous or respiratory administration of rGSN at higher, more frequent doses may allow for improved delivery of rGSN to the lung. Additionally, while rGSN has been shown to bind to lipopolysaccharide (LPS) and improve bacterial killing in some animal models of infection, it is possible that the effects on *L. pneumophila* may differ, due to its intracellular life cycle or atypical LPS structure (Bucki et al., 2005; Piktel et al., 2020; Zaehringer et al., 1995). Legionnaires’ disease is often a difficult and neglected diagnosis, hence its classification as an atypical pneumonia (Mulazimoglu & Yu, 2001; Shimada et al., 2009; Vergis et al., 2000). Broad-spectrum host-directed therapeutics may provide an option for immediate treatment of such severe pneumonias when diagnosis is pending. Administering rGSN alongside antibiotics may be effective in limiting organ damage and disease progression while improving recovery.

Plasma gelsolin is abundant in the blood (Smith DB, Janmey PA, Herbert TJ, 1987). During an infection or injury, pGSN is consumed at the site of insult where it rapidly clears inflammatory and cytotoxic molecules, primarily filamentous actin released from dead cells, resulting in a measurable decrease in circulating pGSN (Bucki et al., 2005; Haddad et al., 1990; Osborn et al., 2008). In several clinical studies examining patients with a wide variety of diseases and injuries, pGSN has been shown to be decreased in diseased patients compared to healthy individuals (Abers et al., 2021; Huang et al., 2011; P. S. Lee et al., 2008; Lind SE, Smith DB, Janmey PA, 1988; Overmyer et al., 2021). The extent of the decrease can exist on a spectrum, where critical disease severity correlates with even further reduction in pGSN than mild disease (Overmyer et al., 2021). Here we show that mice infected with *L. pneumophila* have decreased pGSN concentrations in the serum, corresponding with an increase in the lungs, highlighting its potential as a biomarker for Legionnaires’ disease severity. It remains to be determined if individuals with low pGSN concentrations at baseline are more susceptible to increased disease severity, as pGSN concentration can vary by demographic (Noren Hooten et al., 2023). However, our results suggest that complete absence of gelsolin (and therefore pGSN) In *gsn*^-/-^ mice leaves them at significantly higher risk of developing severe Legionnaires’ disease and succumbing to infection. Even the addition of rGSN at relatively low levels is enough to partially rescue the knockout mice, suggesting pGSN and cGSN play distinct roles in host defense against *L. pneumophila* infection.

A clinical trial investigating the efficacy of recombinant gelsolin administration in treating severe COVID-19 patients found mixed results: gelsolin reduced the frequency of adverse events but did not decrease the length of hospital stay (Dinubile et al., 2022). Varying results of such clinical trials may be due to delivery, dose, timeline of treatment, and genetic variability of the subjects. Additionally, a report of rGSN administration in a severe COVID-19 patient suggests that rGSN administration may improve the disease severity and outcome of these patients (Catteeuw & DiNubile, 2021). Although our study corroborates such anecdotal findings with systemic gelsolin delivery, better targeting rGSN to the lungs, such as through aerosolization, may allow gelsolin to be more concentrated in the lungs. This might overcome issues with dosing and dilution in the body, as well as decreasing the time for the protein to reach the lungs, and potential off target effects. However, the systemic effects of rGSN may be decreased due to the localized delivery to the lungs compared to intravenous delivery. We also show direct treatment of macrophages with rGSN results in a decreased inflammatory cytokine signature in the supernatant of *L. pneumophila*-infected macrophages, further cementing the potential of rGSN as a potential treatment for reducing harmful inflammation during infection.

Upon entering the alveolar space, *L. pneumophila* is phagocytosed by alveolar macrophages. Once inside, *L. pneumophila* halts phagosome-lysosome fusion, forming the *Legionella* containing vacuole (LCV) (Horwitz, 1983). *L. pneumophila* replicates in the LCV, but bacterial and host molecules can escape the LCV through the T4SS and trigger host inflammasome activation (Amer et al., 2006a; Casson et al., 2013; Chin et al., 2020). NAIP5 detects cytosolic flagellin and triggers NLRC4 inflammasome assembly, resulting in caspase-1 mediated bacterial restriction (Amer et al., 2006b; Ren et al., 2006). Caspase-4/11 constitutes the non-canonical inflammasome and mediates cytokine release and pyroptosis once activated (Casson et al., 2013; Cerqueira et al., 2015). However, the substrate repertoire of caspase-4/11 is more diverse than that of other inflammatory caspases and continues to grow as more downstream effectors are discovered (Abu Khweek & Amer, 2020). We have previously shown that caspase-11 modulates actin dynamics during *L. pneumophila* infection through *Limk*-dependent activation of cofilin, leading to phagosome-lysosome fusion and bacterial clearance (Akhter et al., 2012a; Caution et al., 2015a). Here we show that gelsolin is a possible substrate of caspase-4/11 due to its susceptibility to being cleaved by caspase-4 protein. Importantly, this interaction is unique compared to the interaction of gelsolin and caspase-3. Whether caspase-4/11 cleaves gelsolin during *L. pneumophila* infection, and what the identity and function of the caspase-4/11 generated gelsolin products are, remains to be determined.

Mitochondria are the cellular organelles primarily responsible for generating energy for the cell. They are also involved in numerous other cellular processes during infection, including signal transduction, cell death, immune responses, and pathogen defense (Arnoult et al., 2009). During *L. pneumophila* infection, mitochondria produce ROS that contribute to macrophage antibacterial responses (Harada et al., 2007; Kajiwara et al., 2018). *L. pneumophila* also secrete effector proteins that can hijack the mitochondria to manipulate the energetic state of the host cell, allowing the bacteria to survive (Escoll et al., 2017; Fu et al., 2022). Gelsolin binds to voltage dependent anion channel, a mitochondrial pore protein important for regulating mitochondrial membrane permeability, signal transduction, and cell death (García-Bartolomé et al., 2017; Kusano et al., 2000). Our data suggests that gelsolin is important for maintaining mitochondrial function and respiratory potential in macrophages and neutrophils during *L. pneumophila* infection.

Neutrophils are highly abundant granulocytes in the blood and are the first infiltrating cells to arrive at the *L. pneumophila* infected lung (Tateda, Moore, Deng, et al., 2001; Tateda, Moore, Newstead, et al., 2001). Upon stimulation, neutrophils can engulf and kill *L. pneumophila* by generating a ROS burst or eject NETs to trap and kill extracellular bacteria (Koutantou et al., 2025; Price et al., 2021; Ziltener et al., 2016). Although neutrophils are important for pathogen defense, they can cause pathogenic inflammation if they are not properly regulated or resolved after infection (Mayadas et al., 2014). Our data suggests that gelsolin deficiency causes neutrophils to accumulate in the lungs during *L. pneumophila* infection, which correlates with worsened lung pathology. Interestingly, when exposed to *L. pneumophila ex vivo*, gelsolin deficient neutrophils struggle to produce NETs. This may suggest a failure of neutrophils to react to the pathogen, in agreement with our seahorse data, that causes pathogenic inflammation to persist instead of resolve.

Identifying substrates and pathways downstream of non-canonical inflammasome activation will help elucidate its functions and give insight as to how to modulate it to treat diseases. We show that caspase-4 can cleave gelsolin, creating a unique cleavage pattern compared to that of caspase-3 cleaved gelsolin, yet similar to the cleavage products produced by caspase-1. As caspase-3 is an apoptotic executioner caspase, its gelsolin products work to promote apoptosis by collapsing the cytoskeleton and releasing DNA (Geng et al., 1998; Kothakota et al., 1997a). Due to the difference in gelsolin cleavage signatures between caspase-4 and caspase-3, and the ability of caspase-4 to cleave caspase-3 generated gelsolin products, it is possible that caspase-4 can oppose caspase-3 induced apoptosis by creating gelsolin derivatives that are nonfunctional or not apoptotic. Interestingly, caspase-1 cleaved gelsolin produces similar bands to those produced by caspase-4 cleavage. This is likely due to the similarity between caspase-1 and caspase-4 in their tetrapeptide motifs and substrates.

Together, this study shows for the first time that gelsolin is implicated in the response to *L. pneumophila* infection on a cellular and whole organism level, and that rGSN can improve Legionnaires’ disease pathology.

## Materials and Methods

### Bacteria

*Legionella pneumophila* strain LP02 was cultured as previously described (Caution et al., 2015). Briefly, LP02 from frozen glycerol stock was cultured buffered charcoal yeast extract agar and incubated at 37°C for 3 days. Infectious LP02 was cultured in buffered yeast extract containing L-cysteine, ferric nitrate, and thymidine (BYE+) at 37°C and 200RPM for 20 hours then sub-cultured in fresh BYE+ media for 20 hours. Cultures were verified for the presence of motile bacteria. Bacterial concentration was determined by measuring the optical density at 600nm. For animal experiments, infectious inoculum was prepared in sterile PBS to a concentration of 10^7^/mL or 5x10^7^/mL for physiological or severe infection dose infections respectively.

### Mice

C57BL/6 wild type mice were obtained from The Jackson Laboratory. *Gsn*^-/-^ mice were generously provided by Dr. Maya Mikami at Columbia University, New York City, NY. Mice were co-housed for several months before the performance of any experiment. Infections were performed on isoflurane-anesthetized mice by intratracheal instillation of 100uL infectious LP02 inoculum in sterile PBS as previously described (Akhter et al., 2012; Amer et al., 2006; Caution et al., 2015). All mice were housed in a pathogen-free facility, and experiments were conducted with approval from the Animal Care and Use Committee at OSU (Columbus, OH), which is accredited by the American Association for Accreditation of

Laboratory Animal Care International according to guidelines of the Public Health Service as issued in the Guide for the Care and Use of Laboratory Animals.

### Bone marrow macrophage derivation

Macrophages were derived from the bone marrow of mice as previously described (Krause et al., 2018). Bone marrow was flushed from the legs of mice with IMDM supplemented with 50% L929 cell conditioned media, 10% heat-inactivated fetal bovine serum (FBS) (Gibco), and non-essential amino acids (Gibco) (deemed L-cell conditioned media; LCCM) with 0.1% penicillin + streptomycin (LCCM+) and resuspended with a 23g needle to create a single cell suspension. The cell suspension was divided equally into two 145x20mm cell culture plates (Greiner) incubated at 37°C and 5% CO_2_ for six days, with addition of 10mL LCCM+ after three days. Cells were washed twice with PBS (Gibco) and transferred to a new tube. The cell suspension was pelleted and stored at -80°C in recovery cell freezing medium (Gibco) or seeded directly into tissue culture plates (Falcon) for experiments.

### Macrophage cell culture

Macrophages were seeded in LCCM immediately after harvesting or thawed from frozen and incubated at 37°C and 5% CO_2_ for 1-2 days prior to experimentation. Infectious media was prepared in IMDM supplemented with 10% FBS. Macrophages were infected with LP02 at MOI of 0.025 for CFU assays and MOI of 0.5 for all other assays unless otherwise noted. Infectious inoculum was aspirated after 1h to normalize bacterial uptake and replaced with fresh IMDM + 10% FBS for the duration of the infection.

### Recombinant human gelsolin

Recombinant human gelsolin was custom synthesized from Genscript. The amino acid sequence for human gelsolin was acquired from uniprot (ID: P06396-1). Mice were administered 350µg of rGSN in PBS, unless otherwise specified, or same volume of PBS by intraperitoneal injection. Recombinant gelsolin was used in cell culture at a concentration of 100µg/mL in IMDM + 10% FBS unless otherwise specified. Untreated cells received a matching volume of PBS in IMDM + 10% FBS.

### Whole body plethysmography

The lung function of mice was measured using the Finepointe unrestrained whole body plethysmography platform (Buxco). Mice were acclimated in operating chambers for 5 minutes on the two days prior to infection. Baseline (day 0) recordings were taken immediately before infection with 5 minutes of acclimation followed by 5 minutes of recording, with one measurement every 10 seconds using COPD study parameters. Measurements on subsequent days were made at the same time of day and prior to any other manipulations to maintain consistency.

### IHC and histology

Mouse lungs were inflated and fixed with 4% paraformaldehyde in PBS for 48 hours then transferred to 70% EtOH. Embedding, immunohistochemical staining, histological staining, and pathology scoring were performed by Histowiz on the right middle lung lobe from each mouse.

### Lung and spleen homogenization

Left lung lobes and spleens were excised and placed in 1mL sterile PBS + HALT protease inhibitor (Thermofisher) in 2mL Precellys bead tubes (Bertin) on ice before homogenization with a Precellys bead homogenizer (Bertin).

### Determination of *in vivo* bacterial load

Lung or spleen homogenates were serially diluted in sterile PBS and spread onto BCYE agar plates. Plates were incubated at 37°C for 72h before colony forming units were enumerated and normalized to lung weight.

### Serum preparation

Mouse blood was collected via cardiac puncture immediately following CO_2_ euthanasia and was allowed to coagulate for 30 minutes at room temperature. Blood was centrifuged at 2000xg for 10 minutes at 4°C and serum stored at -80°C.

### ELISA

Lung homogenates were centrifuged and the supernatant was stored at -80°C. Cytokine/chemokine ELISAs were performed on lung homogenates or serum using Mesoscale Discovery proinflammatory and cytokine 19-plex multiplex ELISA according to the manufacturer’s instructions (Mesoscale Discovery). Mouse gelsolin (CUSAbio) and human gelsolin (Abcam) were quantified in the lung homogenate and serum according to manufacturer’s instructions. Cell culture supernatants were centrifuged for 10 minutes at 20,000xg and 4°C. IL-1β (DY401), KC/CXCL1 (DY453), IL-6 (DY406), and TNFα (DY410), were quantified by duoset ELISA (R&D) according to the manufacturer’s instructions.

### Single-cell suspension and flow cytometry

CD45-PE conjugated antibody (BD bioscience) was injected retro-orbitally 10 minutes prior to euthanasia. Lungs were dissected into single lobes before being dissociated into single-cell suspension using gentleMACS Octo Dissociator (Miltenyi) and lung dissociation kit according to manufacturer’s instructions (Miltenyi Biotec, 130–095-927). Red blood cells were lysed with 2 mL ACK RBC lysis buffer for 3 min at room temperature. Cells were washed in PBS containing 1% FBS. The single-cell suspension was centrifuged, and the cell pellets were washed twice with PBS. Cell pellets were suspended in 0.5 mL PBS 1% FBS, and the single-cell suspensions were stained with fluorophore-conjugated antibodies against CD11b (), Ly6G (), CD11c (), CD86 (), MHCII (), F4/80 (), CD88 () for fluorometric analysis. Fluorophore-conjugated antibodies were purchased from BD bioscience. Cell suspensions were washed with PBS, resuspended with 1% FBS, and analyzed using a Cytek aurora cytometer.

### Neutrophil isolation

Bone marrow was collected from WT and *gsn*^-/-^ mice and neutrophils were isolated by using the EasySep mouse neutrophil enrichment kit (STEM Cell Technologies, 19762A) following the manufacturer’s instructions.

### Neutrophil culture

Neutrophils were seeded in a poly-L-Lysine-coated 24 well tissue culture plate at 10^6^ per well and allowed to attach for 1h at 37°C and 5% CO_2_. LP02 was added in IMDM + 10% FBS at an MOI of 1.0 for 1h and supernatants were prepared for ELISA.

### NET formation assay

Neutrophils were seeded 2x10^5^/well in a 24-well plate on a poly-L-Lysine-coated glass coverslip. Neutrophils were stimulated for 4h with 100 nM PMA (SigmaAldrich, #P8139) or LP02 at MOI 1 or 10. The cells were fixed with 4% paraformaldehyde, permeabilized with 0.2% Triton X-100 for 10 min, and blocked with 10% goat serum for 30 min at RT. For the visualization of NETs, neutrophils were stained with mouse anti–double-stranded DNA (dsDNA) (Abcam, #ab27156), nuclear envelope, goat anti-rabbit immunoglobulin (Ig)G Alexa Fluor 555 (Thermo Fisher Scientific, #A32732), goat anti-mouse IgG Alexa Fluor (Abcam, #ab150113), and wheat germ agglutinin Alexa Fluor 350 (Thermo Fisher Scientific, #W11263). The coverslips were mounted with Fluoroshield Mounting Medium (Abcam, #ab104135). The cells were visualized by confocal microscopy using a Nikon AX/R confocal microscope. The percentage of cells producing NETs was calculated per field of view. The cells producing NETs were counted when DNA was decondensed and ejected from the cells: percent NETs = (neutrophils with DNA projections × 100)/total number of neutrophils.

### Seahorse assay

XFe96 sensor cartridges (Agilent) were hydrated overnight with seahorse XF calibrant (Agilent) at 37°C and 0% CO_2_. Neutrophils were seeded at 4x10^5^ per well into a poly-L-lysine coated XFe96 cell culture plates (Agilent) and allowed to attach for 1h at 37°C and 5% CO_2_. Neutrophils were treated with fresh media or infected with *L. pneumophila* at an MOI of 0.1 or 1.0 for 1h at 37°C and 5% CO_2_. Macrophages were seeded at 10^5^ per well for 48h then infected with an MOI of 0.5 using the above conditions. After 1h, the media was replaced with complete seahorse media (DMEM + 25mM glucose, 1mM glutamine, 5.6mM pyruvate) and cells were incubated at 37°C in the absence of CO_2_ for 1h prior to the assay. Mitochondrial function was assessed with the Seahorse XF Cell Mito Stress Test Kit (Agilent) using the Seahorse XFe96 Analyzer (Agilent). Drugs were prepared in complete seahorse media as follows: 2µM Oligomycin, 2.5µM FCCP, 0.5µM Rotenone/Antimycin A. Macrophages were counted by staining with Hoechst 33342 (1:4000) for 15 minutes and imaging with a Nikon Ti2 microscope using Nikon NIS-Elements software. Their spare respiratory capacity was normalized to the cell count.

### Transmission electron microscopy

WT and *gsn*^-/-^ macrophages were infected with LP02 MOI 0.5 for 1h, washed with PBS, and fixed in 2.5% glutaraldehyde in 0.1 M phosphate buffer for 2h. Cells were pelleted and prepared for TEM. Samples were imaged with a Tecnai Spirit BioTWIN TEM.

### Immunofluorescence and confocal imaging

Cells on cover slips were fixed with 4% PFA and permeabilized with MeOH then blocked with 0.1% BSA in PBS. Samples were probed with antibodies against TOMM20 (1:300; Sigma WH0009804M1) and *L. pneumophila* (1:300; Invitrogen PA1-7227) in blocking buffer overnight at 4°C. Samples were washed with 0.01% triton X100 in PBS and probed with Alexafluor anti-mouse 594 (1:3000; Invitrogen A-11032) and Alexafluor anti-rabbit 488 (1:3000; Invitrogen A-11008) in blocking buffer for 2h at RT, then DAPI (1:4000) for 15’ at RT. Samples were washed with PBS and H_2_O then mounted onto microscope slides with prolong gold antifade mounting reagent (Invitrogen P36930). Images were captured using an Olympus FV10i confocal microscope with 60x objective and 405nm, 473nm, and 559nm lasers. Image analysis and processing were performed in Olympus Fluoview software.

### Structured illumination microscopy

Samples were prepared for immunofluorescence as described above then imaged, deconvolved, and reconstructed with a Nikon Super Resolution Microscope. Image post processing and video creation was performed using Nikon NIS-Elements.

### Caspase cleavage of gelsolin protein

Recombinant human gelsolin protein was incubated with active recombinant caspase-4 (Enzo Life Sciences), human caspase-3 (Enzo Life Sciences), or human caspase-1 (Enzo Life Sciences) in caspase buffer (Enzo Life Sciences) for 2h at 37°C with gentle agitation. The reaction was halted by addition of protease inhibitor (Thermofisher; I3911) and incubation on ice.

### Western blot

Recombinant protein reactions were boiled in Lamelli buffer at 95°C for 5 minutes, separated by SDS-PAGE, and transferred to a PVDF membrane. Membranes were blocked with 5% nonfat milk in TBST for 1 hour before probing with anti-gelsolin 1:1000 (Abcam 74420) in blocking buffer at 4°C overnight, followed by anti-rabbit IgG (Cell Signaling Technologies) 1:3000 in blocking buffer at room temperature for 2h. Proteins were detected by a luminol based chemiluminescence developing system. **Statistical analysis**. All experiments were performed with at least three independent replicates as indicated in the figure legends. All figures display mean and standard deviation (SD) or standard error of the mean (SEM) from independent experiments. Survival analysis was conducted using Log-rank test and Kaplan-Meier survival curves were used to display the results. Other comparisons between groups were conducted with either two-sample t-test or one-way ANOVA followed by Tukey’s adjustment for multiple comparisons. P < 0.05 was considered statistically significant. All analyses were performed using GraphPad Prism 11.0.2.

## Acknowledgements

We acknowledge OSU CMIF for providing microscopy services for this work. This work was supported by NIH grants P01AI175399, R01AI159452, R01AG082113. The authors declare that they have no conflict of interest.

---

**Figure S1.**
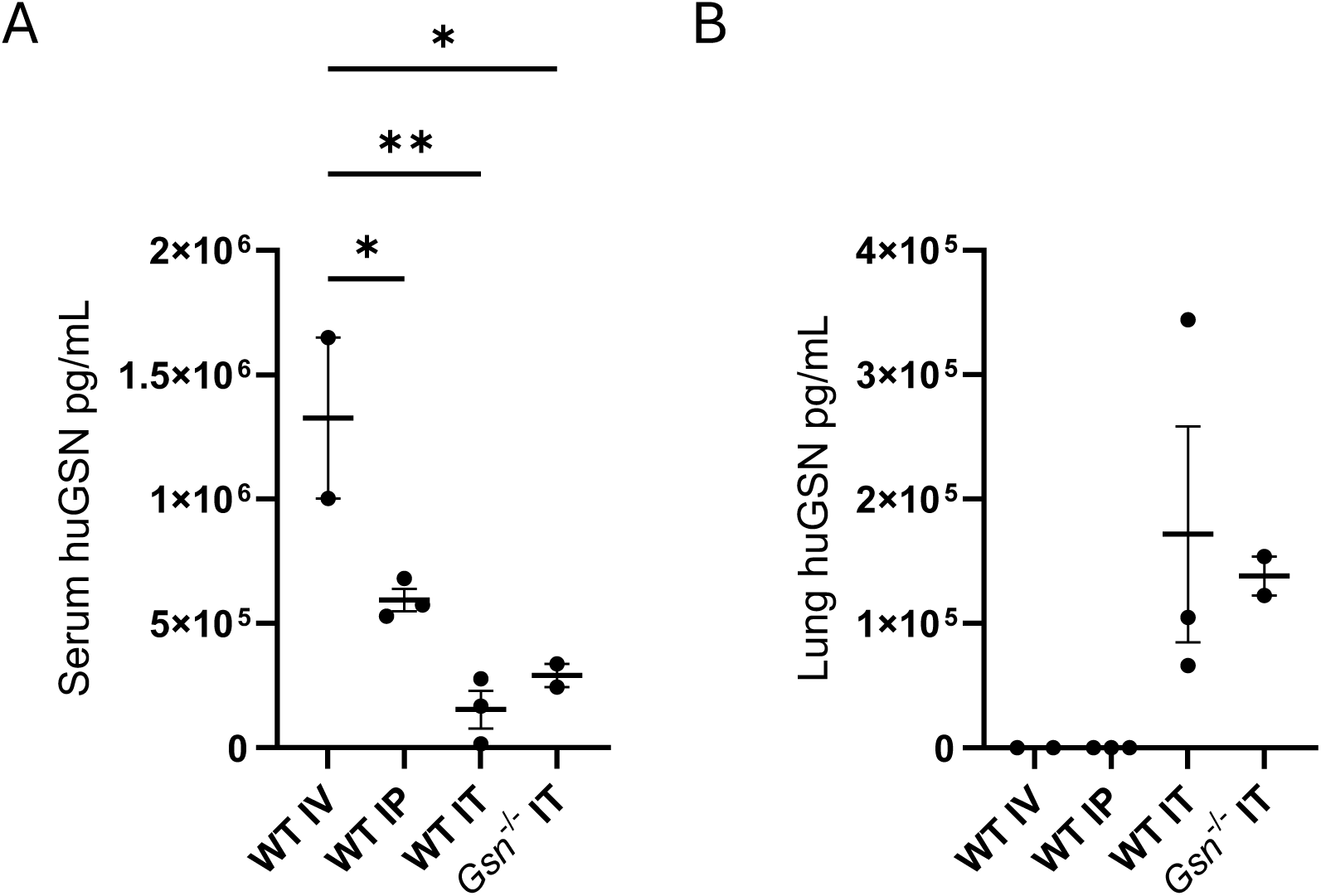
Route of administration determines efficacy and localization of rGSN delivery. Human gelsolin measured by ELISA on serum (A) and lung homogenate (B) of WT or *gsn^-^*^/-^ mice receiving 350µg of rGSN by intravenous (IV) (n=2) or intraperitoneal injection (IP) (n=3), or 180µg of rGSN by intratracheal instillation (IT) (n=2-3). One-way Anova with multiple comparisons. Error bars represent SEM. *p<0.05, **p<0.01.

**Supplemental movie S1.** Reconstructed volume movies from SIM images of WT macrophages infected with LP02 MOI 0.5 for 1h stained with DAPI (cyan), anti-*L. pneumophila* (green), and anti-TOMM20 (red), highlighting the connected mitochondrial morphology in WT macrophages.

**Supplemental movie S2.** Reconstructed volume movies from SIM images of *gsn*^-/-^ macrophages infected with LP02 MOI 0.5 for 1h stained with DAPI (cyan), anti-*L. pneumophila* (green), and anti-TOMM20 (red), highlighting the disrupted mitochondrial morphology in *gsn*^-/-^ macrophages.

## References

Abers, M. S., Delmonte, O. M., Ricotta, E. E., Fintzi, J., Fink, D. L., Almeida de Jesus, A. A., Zarember, K. A., Alehashemi, S., Oikonomou, V., Desai, J. V., Canna, S. W., Shakoory, B., Dobbs, K., Imberti, L., Sottini, A., Quiros-Roldan, E., Castelli, F., Rossi, C., Brugnoni, D., … Notarangelo, L. D. (2021). An immune-based biomarker signature is associated with mortality in COVID-19 patients. JCI Insight, 6(1), 1–20. 10.1172/jci.insight.144455

Abu Khweek, A., & Amer, A. O. (2020). Pyroptotic and non-pyroptotic effector functions of caspase-11. Immunological Reviews, 297(1), 39–52. 10.1111/imr.12910

Akhter, A., Caution, K., Abu Khweek, A., Tazi, M., Abdulrahman, B. A., Abdelaziz, D. H. A., Voss, O. H., Doseff, A. I., Hassan, H., Azad, A. K., Schlesinger, L. S., Wewers, M. D., Gavrilin, M. A., & Amer, A. O. (2012a). Caspase-11 Promotes the Fusion of Phagosomes Harboring Pathogenic Bacteria with Lysosomes by Modulating Actin Polymerization. Immunity, 37(1), 35–47. 10.1016/j.immuni.2012.05.001

Akhter, A., Caution, K., Abu Khweek, A., Tazi, M., Abdulrahman, B. A., Abdelaziz, D. H. A., Voss, O. H., Doseff, A. I., Hassan, H., Azad, A. K., Schlesinger, L. S., Wewers, M. D., Gavrilin, M. A., & Amer, A. O. (2012b). Caspase-11 Promotes the Fusion of Phagosomes Harboring Pathogenic Bacteria with Lysosomes by Modulating Actin Polymerization. Immunity, 37(1), 35–47. 10.1016/j.immuni.2012.05.001

Amer, A., Franchi, L., Kanneganti, T. D., Body-Malapel, M., Özören, N., Brady, G., Meshinchi, S., Jagirdar, R., Gewirtz, A., Akira, S., & Núñez, G. (2006a). Regulation of Legionella phagosome maturation and infection through flagellin and host Ipaf. Journal of Biological Chemistry, 281(46), 35217–35223. 10.1074/jbc.M604933200

Amer, A., Franchi, L., Kanneganti, T. D., Body-Malapel, M., Özören, N., Brady, G., Meshinchi, S., Jagirdar, R., Gewirtz, A., Akira, S., & Núñez, G. (2006b). Regulation of Legionella phagosome maturation and infection through flagellin and host Ipaf. Journal of Biological Chemistry, 281(46), 35217–35223. 10.1074/jbc.M604933200

Arnoult, D., Carneiro, L., Tattoli, I., & Girardin, S. E. (2009). The role of mitochondria in cellular defense against microbial infection. Seminars in Immunology, 21(4), 223–232. 10.1016/j.smim.2009.05.009

Bryan, J. (1988). Gelsolin has three actin-binding sites. Journal of Cell Biology, 106(5), 1553–1562. 10.1083/jcb.106.5.1553

Bucki, R., Byfield, F. J., Kulakowska, A., Mccormick, M. E., Drozdowski, W., Namiot, Z., Hartung, T., & Janmey, P. A. (2014). Extracellular Gelsolin Binds Lipoteichoic Acid and Modulates Components 1. 181(2008), 4936–4944.

Bucki, R., Georges, P. C., Espinassous, Q., Funaki, M., Pastore, J. J., Chaby, R., & Janmey, P. A. (2005). Inactivation of endotoxin by human plasma gelsolin. Biochemistry, 44(28), 9590–9597. 10.1021/bi0503504

Bucki, R., Kułakowska, A., Byfield, F. J., Zendzian-Piotrowska, M., Baranowski, M., Marzec, M., Winer, J. P., Ciccarelli, N. J., Górski, J., Drozdowski, W., Bittman, R., & Janmey, P. A. (2010). Plasma gelsolin modulates cellular response to sphingosine 1-phosphate. American Journal of Physiology - Cell Physiology, 299(6), 1516–1523. 10.1152/ajpcell.00051.2010

Casson, C. N., Copenhaver, A. M., Zwack, E. E., Nguyen, H. T., Strowig, T., Javdan, B., Bradley, W. P., Fung, T. C., Flavell, R. A., Brodsky, I. E., & Shin, S. (2013). Caspase-11 Activation in Response to Bacterial Secretion Systems that Access the Host Cytosol. PLoS Pathogens, 9(6). 10.1371/journal.ppat.1003400

Catteeuw, J. V., & DiNubile, M. J. (2021). Recombinant human plasma gelsolin (rhu-pGSN) in a patient hospitalized with critical COVID-19 pneumonia. Clinical Infection in Practice, 12(January). 10.1016/j.clinpr.2021.100088

Caution, K., Gavrilin, M. A., Tazi, M., Kanneganti, A., Layman, D., Hoque, S., Krause, K., & Amer, A. O. (2015a). Caspase-11 and caspase-1 differentially modulate actin polymerization via RhoA and Slingshot proteins to promote bacterial clearance. Scientific Reports, 5(November), 1–13. 10.1038/srep18479

Caution, K., Gavrilin, M. A., Tazi, M., Kanneganti, A., Layman, D., Hoque, S., Krause, K., & Amer, A. O. (2015b). Caspase-11 and caspase-1 differentially modulate actin polymerization via RhoA and Slingshot proteins to promote bacterial clearance. Scientific Reports, 5(November), 1–13. 10.1038/srep18479

Cerqueira, D. M., Pereira, M. S. F., Silva, A. L. N., Cunha, L. D., & Zamboni, D. S. (2015). Caspase-1 but Not Caspase-11 Is Required for NLRC4-Mediated Pyroptosis and Restriction of Infection by Flagellated Legionella Species in Mouse Macrophages and In Vivo. The Journal of Immunology, 195(5), 2303–2311. 10.4049/jimmunol.1501223

Cheng, Y., Hu, X., Liu, C., Chen, M., Wang, J., Wang, M., Gao, F., Han, J., Sun, D., Zhang, C., & Min, R. (2017). Gelsolin inhibits the inflammatory process induced by LPS. Cellular Physiology and Biochemistry, 41(1), 205–212. 10.1159/000456043

Chin, E. N., Yu, C., Vartabedian, V. F., Jia, Y., Kumar, M., Gamo, A. M., Vernier, W., Ali, S. H., Kissai, M., Lazar, D. C., Nguyen, N., Pereira, L. E., Benish, B., Woods, A. K., Joseph, S. B., Chu, A., Johnson, K. A., Sander, P. N., Martínez-peña, F., … Lairson, L. L. (2020). HO. 999(August), 993–999.

Christofidou-Solomidou, M., Scherpereel, A., Solomides, C. C., Christie, J. O., Stossel, T. P., Goelz, S., & DiNubile, M. J. (2002). Recombinant plasma gelsolin diminishes the acute inflammatory response to hyperoxia in mice. Journal of Investigative Medicine, 50(1), 54–60. 10.2310/6650.2002.33518

Cui, T. X., Brady, A. E., Zhang, Y.-J., Fulton, C. T., & Popova, A. P. (2022). Gelsolin Attenuates Neonatal Hyperoxia-Induced Inflammatory Responses to Rhinovirus Infection and Preserves Alveolarization. Frontiers in Immunology, 13(January), 1–13. 10.3389/fimmu.2022.792716

Dartevel, A., Galerneau, L.-M., Peigne, V., Sedillot, N., Ehrmann, S., Lautrette, A., Klouche, K., Poissy, J., Thiery, G., Sauneuf, B., Rigaud, J.-P., Ramakers, M., Daubin, C., Schwebel, C., & Terzi, N. (2025). Clinical features and prognosis of severe legionnaires’ disease requiring intensive care unit admission: a multicentric retrospective cohort study. Pneumonia, 17(1). 10.1186/s41479-025-00173-z

De Filippis, P., Mozzetti, C., Messina, A., & D’Alò, G. L. (2018). Prevalence of Legionella in retirement homes and group homes water distribution systems. Science of the Total Environment, 643, 715–724. 10.1016/j.scitotenv.2018.06.216

DiNubile, M. J., Levinson, S. L., Stossel, T. P., Lawrenz, M. B., & Warawa, J. M. (2020). Recombinant human plasma gelsolin improves survival and attenuates lung injury in a murine model of multidrug-resistant Pseudomonas aeruginosa pneumonia. Open Forum Infectious Diseases, 7(8), 1–8. 10.1093/ofid/ofaa236

Dinubile, M. J., Parra, S., Salomó, A. C., & Levinson, S. L. (2022). Adjunctive Recombinant Human Plasma Gelsolin for Severe Coronavirus Disease 2019 Pneumonia. Open Forum Infectious Diseases, 0(0), 1–8. 10.1093/ofid/ofac357

DiNubile, M. J., Stossel, T. P., Ljunghusen, O. C., Ferrara, J. L. M., & Antin, J. H. (2002). Prognostic implications of declining plasma gelsolin levels after allogeneic stem cell transplantation. Blood, 100(13), 4367–4371. 10.1182/blood-2002-06-1672

Douda, D. N., Khan, M. A., Grasemann, H., & Palaniyar, N. (2015). SK3 channel and mitochondrial ROS mediate NADPH oxidase-independent NETosis induced by calcium influx. Proceedings of the National Academy of Sciences of the United States of America, 112(9), 2817–2822. 10.1073/pnas.1414055112

Esawy, M. M., Makram, W. K., Albalat, W., & Shabana, M. A. (2020). Plasma gelsolin levels in patients with psoriatic arthritis: a possible novel marker. Clinical Rheumatology, 39(6), 1881–1888. 10.1007/s10067-020-04959-y

Escoll, P., Song, O. R., Viana, F., Steiner, B., Lagache, T., Olivo-Marin, J. C., Impens, F., Brodin, P., Hilbi, H., & Buchrieser, C. (2017). Legionella pneumophila Modulates Mitochondrial Dynamics to Trigger Metabolic Repurposing of Infected Macrophages. Cell Host and Microbe, 22(3), 302–316.e7. 10.1016/j.chom.2017.07.020

Fang, Z., Zhou, X., Liao, H., & Xu, H. (2023). A meta-analysis of Legionella pneumophila contamination in hospital water systems. American Journal of Infection Control, 51(11), 1250–1262. 10.1016/j.ajic.2023.04.002

Fettucciari, K., Ponsini, P., Palumbo, C., Rosati, E., Mannucci, R., Bianchini, R., Modesti, A., & Marconi, P. (2015). Macrophage induced gelsolin in response to Group B Streptococcus (GBS) infection. Cellular Microbiology, 17(1), 79–104. 10.1111/cmi.12338

Fu, J., Zhou, M., Gritsenko, M. A., Nakayasu, E. S., Song, L., & Luo, Z. Q. (2022). Legionella pneumophila modulates host energy metabolism by ADP-ribosylation of ADP/ATP translocases. ELife, 11(Lcv), 1–20. 10.7554/eLife.73611

García-Bartolomé, A., Peñas, A., Marín-Buera, L., Lobo-Jarne, T., Pérez-Pérez, R., Morán, M., Arenas, J., Martín, M. A., & Ugalde, C. (2017). Respiratory chain enzyme deficiency induces mitochondrial location of actin-binding gelsolin to modulate the oligomerization of VDAC complexes and cell survival. Human Molecular Genetics, 26(13), 2493–2506. 10.1093/hmg/ddx144

Geng, Y. J., Azuma, T., Tang, J. X., Hartwig, J. H., Muszynski, M., Wu, Q., Libby, P., & Kwiatkowski, D. J. (1998). Caspase-3-induced gelsolin fragmentation contributes to actin cytoskeletal collapse, nucleolysis, and apoptosis of vascular smooth muscle cells exposed to proinflammatory cytokines. European Journal of Cell Biology, 77(4), 294–302. 10.1016/S0171-9335(98)80088-5

Haddad, J. G., Harper, K. D., Guoth, M., Pietra, G. G., & Sanger, J. W. (1990). Angiopathic consequences of saturating the plasma scavenger system for actin. Proceedings of the National Academy of Sciences of the United States of America, 87(4), 1381–1385. 10.1073/pnas.87.4.1381

Harada, T., Miyake, M., & Imai, Y. (2007). Evasion of Legionella pneumophila from the bactericidal system by reactive oxygen species (ROS) in macrophages. Microbiology and Immunology, 51(12), 1161–1170. 10.1111/j.1348-0421.2007.tb04011.x

Horwitz, M. A. (1983). The legionnaires’ disease bacterium (legionella pneumophila) inhibits phagosome-lysosome fusion in human monocytes. Journal of Experimental Medicine, 158(6), 2108–2126. 10.1084/jem.158.6.2108

Huang, L. F., Yao, Y. M., Li, J. F., Dong, N., Liu, C., Yu, Y., He, L. X., & Sheng, Z. Y. (2011). Reduction of plasma gelsolin levels correlates with development of multiple organ dysfunction syndrome and fatal outcome in burn patients. PLoS ONE, 6(10). 10.1371/journal.pone.0025748

Janmey, P. A., Chaponnier, C., Lind, S. E., Zaner, K. S., Stossel, T. P., & Yin, H. L. (1985). Interactions of Gelsolin and Gelsolin-Actin Complexes with Actin. Effects of Calcium on Actin Nucleation, Filament Severing, and End Blocking. Biochemistry, 24(14), 3714–3723. 10.1021/bi00335a046

Kajiwara, C., Kusaka, Y., Kimura, S., Yamaguchi, T., Nanjo, Y., Ishii, Y., Udono, H., Standiford, T. J., & Tateda, K. (2018). Metformin Mediates Protection against Legionella Pneumonia through Activation of AMPK and Mitochondrial Reactive Oxygen Species. The Journal of Immunology, 200(2), 623–631. 10.4049/jimmunol.1700474

Kayagaki, N., Stowe, I. B., Lee, B. L., O’Rourke, K., Anderson, K., Warming, S., Cuellar, T., Haley, B., Roose-Girma, M., Phung, Q. T., Liu, P. S., Lill, J. R., Li, H., Wu, J., Kummerfeld, S., Zhang, J., Lee, W. P., Snipas, S. J., Salvesen, G. S., … Dixit, V. M. (2015). Caspase-11 cleaves gasdermin D for non-canonical inflammasome signalling. Nature, 526(7575), 666–671. 10.1038/nature15541

Kayagaki, N., Warming, S., Lamkanfi, M., Walle, L. Vande, Louie, S., Dong, J., Newton, K., Qu, Y., Liu, J., Heldens, S., Zhang, J., Lee, W. P., Roose-Girma, M., & Dixit, V. M. (2011). Non-canonical inflammasome activation targets caspase-11. Nature, 479(7371), 117–121. 10.1038/nature10558

Kobzik, L., Yang, Z., Bedugnis, A., Levinson, S., DiNubile, M., Stossel, T., & Lu, Q. (2020). Delayed administration of recombinant plasma gelsolin improves survival in a murine model of severe influenza. F1000Research, 8, 1–19. 10.12688/f1000research.21082.2

Kothakota, S., Azuma, T., Reinhard, C., Klippel, A., Tang, J., Chu, K., McGarry, T. J., Kirschner, M. W., Koths, K., Kwiatkowski, D. J., & Williams, L. T. (1997a). Caspase-3-generated fragment of gelsolin: Effector of morphological change in apoptosis. Science, 278(5336), 294–298. 10.1126/science.278.5336.294

Kothakota, S., Azuma, T., Reinhard, C., Klippel, A., Tang, J., Chu, K., McGarry, T. J., Kirschner, M. W., Koths, K., Kwiatkowski, D. J., & Williams, L. T. (1997b). Caspase-3-generated fragment of gelsolin: Effector of morphological change in apoptosis. Science, 278(5336), 294–298. 10.1126/science.278.5336.294

Koutantou, M., Konstantinidis, T., Chochlakis, D., Xingi, E., Psaroulaki, A., Tsiotis, G., Kambas, K., & Angelakis, E. (2025). IL-1beta expressing neutrophil extracellular traps in Legionella pneumophila infection. Frontiers in Immunology, 16(June), 1–11. 10.3389/fimmu.2025.1573151

Koya, R. C., Fujita, H., Shimizu, S., Ohtsu, M., Takimoto, M., Tsujimoto, Y., & Kuzumaki, N. (2000). Gelsolin inhibits apoptosis by blocking mitochondrial membrane potential loss and cytochrome c release. Journal of Biological Chemistry, 275(20), 15343–15349. 10.1074/jbc.275.20.15343

Krause, K., Caution, K., Badr, A., Hamilton, K., Saleh, A., Patel, K., Seveau, S., Hall-Stoodley, L., Hegazi, R., Zhang, X., Gavrilin, M. A., & Amer, A. O. (2018). CASP4/caspase-11 promotes autophagosome formation in response to bacterial infection. Autophagy, 14(11), 1928–1942. 10.1080/15548627.2018.1491494

Krause, K., Daily, K., Estfanous, S., Hamilton, K., Badr, A., Abu Khweek, A., Hegazi, R., Anne, M. N., Klamer, B., Zhang, X., Gavrilin, M. A., Pancholi, V., & Amer, A. O. (2019). Caspase-11 counteracts mitochondrial ROS-mediated clearance of Staphylococcus aureus in macrophages. EMBO Reports, 20(12), 1–17. 10.15252/embr.201948109

Kusano, H., Shimizu, S., Koya, R. C., Fujita, H., Kamada, S., Kuzumaki, N., & Tsujimoto, Y. (2000). Human gelsolin prevents apoptosis by inhibiting apoptotic mitochondrial changes via closing VDAC. Oncogene, 19(42), 4807–4814. 10.1038/sj.onc.1203868

Kwiatkowski, D. J., Mehl, R., Izumo, S., Nadal-Ginard, B., & Yin, H. L. (1988). Muscle is the major source of plasma gelsolin. Journal of Biological Chemistry, 263(17), 8239–8243. 10.1016/s0021-9258(18)68469-8

Lee, J., Sasaki, F., Koike, E., Cho, M., Lee, Y., Dho, S. H., Lee, J., Lee, E., Toyohara, E., Sunakawa, M., Ishibashi, M., Hung, H. H., Nishioka, S., Komine, R., Okura, C., Shimizu, M., Ikawa, M., Yoshimura, A., Morita, R., & Kim, L. K. (2024). Gelsolin alleviates rheumatoid arthritis by negatively regulating NLRP3 inflammasome activation. Cell Death and Differentiation, 31(12), 1679–1694. 10.1038/s41418-024-01367-6

Lee, P. S., Drager, L. R., Stossel, T. P., Moore, F. D., & Rogers, S. O. (2006). Relationship of plasma gelsolin levels to outcomes in critically III surgical patients. Annals of Surgery, 243(3), 399–403. 10.1097/01.sla.0000201798.77133.55

Lee, P. S., Patel, S. R., Christiani, D. C., Bajwa, E., Stossel, T. P., & Waxman, A. B. (2008). Plasma gelsolin depletion and circulating actin in sepsis - A pilot study. PLoS ONE, 3(11), 1–5. 10.1371/journal.pone.0003712

Lee, P. S., Waxman, A. B., Cotich, K. L., Chung, S. W., Perrella, M. A., & Stossel, T. P. (2007). Plasma gelsolin is a marker and therapeutic agent in animal sepsis. Critical Care Medicine, 35(3), 849–855. 10.1097/01.CCM.0000253815.26311.24

Lind, S. E., Smith, D. B., Janmey, P. A., & Stossel, T. P. (1986). Role of plasma gelsolin and the vitamin D-binding protein in clearing actin from the circulation. Journal of Clinical Investigation, 78(3), 736–742. 10.1172/JCI112634

Lind SE, Smith DB, Janmey PA, S. TP. (1988). Depression of gelsolin levels and detection of gelsolin-actin complexes in plasma of patients with acute lung injury. Am Rev Respir Dis., 138(2), 429–434.

Luo, Z. Q. (2012). Legionella secreted effectors and innate immune responses. Cellular Microbiology, 14(1), 19–27. 10.1111/j.1462-5822.2011.01713.x

Lupia, T., Corcione, S., Shbaklo, N., Rizzello, B., De Benedetto, I., Concialdi, E., Navazio, A. S., Penna, M., Brusa, M. T., & De Rosa, F. G. (2023). Legionella pneumophila Infections during a 7-Year Retrospective Analysis (2016–2022): Epidemiological, Clinical Features and Outcomes in Patients with Legionnaires’ Disease. Microorganisms, 11(2). 10.3390/microorganisms11020498

Mannherz, H. G., Budde, H., Jarkas, M., Hassoun, R., Malek-Chudzik, N., Mazur, A. J., Skuljec, J., Pul, R., Napirei, M., & Hamdani, N. (2024a). Reorganization of the actin cytoskeleton during the formation of neutrophil extracellular traps (NETs). European Journal of Cell Biology, 103(2), 151407. 10.1016/j.ejcb.2024.151407

Mannherz, H. G., Budde, H., Jarkas, M., Hassoun, R., Malek-Chudzik, N., Mazur, A. J., Skuljec, J., Pul, R., Napirei, M., & Hamdani, N. (2024b). Reorganization of the actin cytoskeleton during the formation of neutrophil extracellular traps (NETs). European Journal of Cell Biology, 103(2), 151407. 10.1016/j.ejcb.2024.151407

Mayadas, T., Cullere, X., & Lowell, C. (2014). The Multifaceted Functions of Neutrophils. Annual Review of Pathology: Mechanisms of Disease, 9, 181–218. 10.1146/annurev-pathol-020712-164023.The

Mulazimoglu, L., & Yu, V. L. (2001). Can Legionnaires disease be diagnosed by clinical criteria? A critical review. Chest, 120(4), 1049–1053. 10.1378/chest.120.4.1049

Noren Hooten, N., Mode, N. A., Kowalik, E., Omoniyi, V., Zonderman, A. B., Ezike, N., DiNubile, M. J., Levinson, S. L., & Evans, M. K. (2023). Plasma gelsolin levels are associated with diabetes, sex, race, and poverty. Journal of Translational Medicine, 21(1), 1–10. 10.1186/s12967-023-04026-5

Osborn, T. M., Verdrengh, M., Stossel, T. P., & Bokarewa, M. (2008). Decreased levels of the gelsolin plasma isoform in patients with rheumatoid arthritis. Arthritis Research and Therapy, 10(5), 1–9. 10.1186/ar2520

Overmyer, K. A., Shishkova, E., Miller, I. J., Balnis, J., Bernstein, M. N., Peters-Clarke, T. M., Meyer, J. G., Quan, Q., Muehlbauer, L. K., Trujillo, E. A., He, Y., Chopra, A., Chieng, H. C., Tiwari, A., Judson, M. A., Paulson, B., Brademan, D. R., Zhu, Y., Serrano, L. R., … Jaitovich, A. (2021). Large-Scale Multi-omic Analysis of COVID-19 Severity. Cell Systems, 12(1), 23–40.e7. 10.1016/j.cels.2020.10.003

Pereira, M. S. F., Marques, G. G., DelLama, J. E., & Zamboni, D. S. (2011). The Nlrc4 inflammasome contributes to restriction of pulmonary infection by flagellated Legionella spp. that trigger pyroptosis. Frontiers in Microbiology, 2(FEB), 1–6. 10.3389/fmicb.2011.00033

Pereira, M. S. F., Morgantetti, G. F., Massis, L. M., Horta, C. V, Hori, J. I., & Zamboni, D. S. (2011). Activation of NLRC4 by Flagellated Bacteria Triggers Caspase-1–Dependent and –Independent Responses To Restrict Legionella pneumophila Replication in Macrophages and In Vivo. The Journal of Immunology, 187(12), 6447–6455. 10.4049/jimmunol.1003784

Piktel, E., Wnorowska, U., Cieśluk, M., Deptuła, P., Prasad, S. V., Król, G., Durnaś, B., Namiot, A., Markiewicz, K. H., Niemirowicz-Laskowska, K., Wilczewska, A. Z., Janmey, P. A., Reszeć, J., & Bucki, R. (2020). Recombinant human plasma gelsolin stimulates phagocytosis while diminishing excessive inflammatory responses in mice with Pseudomonas Aeruginosa Sepsis. International Journal of Molecular Sciences, 21(7). 10.3390/ijms21072551

Price, C. T. D., Hanford, H. E., Vashishta, A., Ozanic, M., Santic, M., & Uriarte, S. (2021). Dot / Icm-Dependent Restriction of Legionella pneumophila within Neutrophils. 12(3), 1–18.

Ren, T., Zamboni, D. S., Roy, C. R., Dietrich, W. F., & Vance, R. E. (2006). Flagellin-deficient Legionella mutants evade caspase-1- and Naip5-mediated macrophage immunity. PLoS Pathogens, 2(3), 0175–0183. 10.1371/journal.ppat.0020018

Self, W. H., Wunderink, R. G., Dinubile, M. J., Stossel, T. P., Levinson, S. L., Williams, D. J., Anderson, E. J., Bramley, A. M., Jain, S., Edwards, K. M., & Grijalva, C. G. (2019). Low Admission Plasma Gelsolin Concentrations Identify Community-acquired Pneumonia Patients at High Risk for Severe Outcomes. Clinical Infectious Diseases, 69(7), 1218–1225. 10.1093/cid/ciy1049

Serrander, L., Skarman, P., Rasmussen, B., Witke, W., Lew, D. P., Krause, K.-H., Stendahl, O., & Nüße, O. (2000). Selective Inhibition of IgG-Mediated Phagocytosis in Gelsolin-Deficient Murine Neutrophils. The Journal of Immunology, 165(5), 2451–2457. 10.4049/jimmunol.165.5.2451

Shi, S. S., Chen, C., Zhao, D. Y., Liu, X. W., Cheng, B. L., Wu, S. J., Lin, R., Tan, L. H., Fang, X. M., & Shu, Q. (2014). The role of plasma gelsolin in cardiopulmonary bypass induced acute lung injury in infants and young children: A pilot study. BMC Anesthesiology, 14(1), 1–9. 10.1186/1471-2253-14-67

Shimada, T., Noguchi, Y., Jackson, J. L., Miyashita, J., Hayashino, Y., Kamiya, T., Yamazaki, S., Matsumura, T., & Fukuhara, S. (2009). Systematic review and metaanalysis: Urinary antigen tests for legionellosis. Chest, 136(6), 1576–1585. 10.1378/chest.08-2602

Smith DB, Janmey PA, Herbert TJ, L. S. (1987). Quantitative measurement of plasma gelsolin and its incorporation into fibrin clots. J Lab Clin Med, 110(2), 89–95.

Stella, M. C., Schauerte, H., Straub, K. L., & Leptin, M. (1994). Identification of secreted and cytosolic gelsolin in Drosophila. Journal of Cell Biology, 125(3), 607–616. 10.1083/jcb.125.3.607

Tateda, K., Moore, T. A., Deng, J. C., Newstead, M. W., Zeng, X., Matsukawa, A., Swanson, M. S., Yamaguchi, K., & Standiford, T. J. (2001). Early Recruitment of Neutrophils Determines Subsequent T1/T2 Host Responses in a Murine Model of Legionella pneumophila Pneumonia. The Journal of Immunology, 166(5), 3355–3361. 10.4049/jimmunol.166.5.3355

Tateda, Kazuhiro., Moore, T. A., Newstead, M. W., Tsai, W. C., Zeng, X., Deng, J. C., Chen, G., Reddy, R., Yamaguchi, K., & Standiford, T. J. (2001). Chemokine-dependent neutrophil recruitment in a murine model of Legionella pneumonia: Potential role of neutrophils as immunoregulatory cells. Infection and Immunity, 69(4), 2017–2024. 10.1128/IAI.69.4.2017-2024.2001

Van Heijnsbergen, E., Schalk, J. A. C., Euser, S. M., Brandsema, P. S., Den Boer, J. W., & De Roda Husman, A. M. (2015). Confirmed and potential sources of Legionella reviewed. Environmental Science and Technology, 49(8), 4797–4815. 10.1021/acs.est.5b00142

Vergis, E., Akbas, E., & Yu, V. L. (2000). Legionella as a Cause of Severe Pneumonia. SEMINARS IN RESPIRATORY AND CRITICAL CARE MEDICINE, 21(4), 295–304. 10.1086/603594

Yang, Z., Chiou, T. T. Y., Stossel, T. P., & Kobzik, L. (2015). Plasma gelsolin improves lung host defense against pneumonia by enhancing macrophage NOS3 function. American Journal of Physiology - Lung Cellular and Molecular Physiology, 309(1), L11–L16. 10.1152/ajplung.00094.2015

Zaehringer, U., Knirel, Y., Lindner, B., & Sonesson, A. (1995). The lipopolysaccaride of Legionella pneumophila serogroup 1 (strain. 1(February).

Zhao, Y., Yang, J., Shi, J., Gong, Y. N., Lu, Q., Xu, H., Liu, L., & Shao, F. (2011). The NLRC4 inflammasome receptors for bacterial flagellin and type III secretion apparatus. Nature, 477(7366), 596–602. 10.1038/nature10510

Ziltener, P., Reinheckel, T., & Oxenius, A. (2016). Neutrophil and Alveolar Macrophage-Mediated Innate Immune Control of Legionella pneumophila Lung Infection via TNF and ROS. PLoS Pathogens, 12(4), 1–25. 10.1371/journal.ppat.1005591

